# Weak-Instrument Robust Tests in Two-Sample Summary-Data Mendelian Randomization

**DOI:** 10.1101/769562

**Authors:** Sheng Wang, Hyunseung Kang

## Abstract

Mendelian randomization (MR) is a popular method in genetic epidemiology to estimate the effect of an exposure on an outcome using genetic variants as instrumental variables (IV), with two-sample summary-data MR being the most popular due to privacy. Unfortunately, many MR methods for two-sample summary data are not robust to weak instruments, a common phenomena with genetic instruments; many of these methods are biased and no existing MR method has Type I error control under weak instruments. In this work, we propose test statistics that are robust to weak instruments by extending Anderson-Rubin, Kleibergen, and conditional likelihood ratio tests in econometrics to the two-sample summary data setting. We conclude with a simulation and an empirical study and show that the proposed tests control size and have better power than current methods.

## 1 Introduction

Recently, Mendelian randomization (MR) is a popular method in genetic epidemiology to study the effect of modifiable exposures on health outcomes by using genetic variants as instrumental variables (IV). In a nutshell, MR finds instruments, typically single nucleotide polymorphisms (SNPs), from publicly available genome-wide association studies (GWAS) and the instruments must be (A1) associated with the exposure; (A2) independent of the unmeasured confounder; and (A3) independent of the outcome variable after conditioning on the exposure (Davey Smith and Ebrahim, 2003; Lawlor et al., 2008) Typically, two non-overlapping GWAS are used to find instruments, one GWAS studying the exposure and another GWAS studying the outcome. Also, due to privacy, when estimating the exposure effect, only summary statistics instead of individual-level data are extracted for analysis. This setup is commonly known as two-sample summary-data MR (Pierce and Burgess, 2013; Burgess et al., 2013, 2015).

The focus of this paper is on testing the exposure effect in two-sample summary-data MR when (A1) is violated, or more precisely when the instruments are weakly associated with the exposure. Many genetic instruments in MR studies only explain a fraction of the variation in the exposure. Using these weak instruments can introduce bias and inflate Type I errors (Burgess and Thompson, 2011). Weak IVs can also amplify bias from minor violations of (A2) and (A3) (Small and Rosenbaum, 2008). Unfortunately, many popular MR methods and software assume instruments are strongly associated with the exposure; many go one step further and assume that the correlation between each instrument and exposure has no sampling error (Bowden et al., 2016b). For example, methods such as the inverse-variance weighted estimator (IVW) (Burgess et al., 2013), MR-Egger regression (Bowden et al., 2015), weighted median estimator (W.Median) (Bowden et al., 2016a) and the modal estimator (Hartwig et al., 2017) typically assume that each instrument’s correlation to the exposure of interest is measured without error. In Section 4, we numerically demonstrate the seriousness of making such assumptions in MR by “stress-testing” these methods’ performance on a real MR data set, akin to an exercise done by Bound et al. (1995) in econometrics for single-sample, individual-level data.

Many works in econometrics have dealt with weak instruments; see Stock et al. (2002) for an overview. In particular, the Anderson-Rubin (AR) test (Anderson et al., 1949), the Kleibergen (K) test (Kleibergen, 2002), and the conditional likelihood ratio (CLR) test (Moreira, 2003) provide Type I error control regardless of instruments’ magnitude of association to the exposure, also called instruments’ strength. More formally, the three methods satisfy the necessary requirement for valid 1 − *α* confidence intervals with weak instruments, where the confidence interval adapts and becomes infinite in the presence of weak IVs to maintain 1 − *α* coverage (Dufour, 1997). However, all these methods assume that individual-level data is available to compute the test statistics and the data comes from the same sample. In GWAS and MR, one rarely has access to individual-level data due to privacy concerns and is forced to work with anonymized summary statistics from multiple GWAS. To the best of our knowledge, no methods in two-sample summary-data MR provide Type I error control when the relationship between the exposure and the instruments is weak, even irrelevant.

Our contribution is to propose weak instrument robust tests for two-sample summary-data MR. We extend the three aforementioned weak-instrument robust tests in econometrics, AR test, the K test, and the CLR test, to work with two-sample summary data by leveraging recent work by Choi et al. (2018) who worked with two-sample, but individual data. We show that under the two-sample summary-data model and weak-IV asymptotics of Staiger and Stock (1997), these modified tests, which we call mrAR, mrK, and mrCLR, asymptotically control Type I error for testing the exposure effect. In the supplementary materials, we also introduce point estimators based on these tests, most notably the MR limited information maximum likelihood estimator (mrLIML) based on taking the minimum of the mrAR test. mrLIML is similar to the original limited information maximum likelihood (LIML) estimator (Anderson et al., 1949) and we show an equivalence relationship between mrLIML and the recent profile-likelihood estimator proposed by Zhao et al. (2018). We conclude with a simulation and a prototypical MR data example concerning the effect of body mass index (BMI) on systolic blood pressure (SBP).

## 2 Setup and Method

### 2.1 Review: Two-Sample Summary Data in MR

We review the data generating model underlying MR. Suppose we have two independent groups of people, with *n*_1_ and *n*_2_ participants each of the two groups. For each individual *i* in sample *l* = 1, 2, let *Y*_*li*_ ∈ ℝ denote his/her outcome, *D*_*li*_ ∈ ℝ denote his/her exposure, and *Z*_*li*_ ∈ ℝ^*L*^ denote his/her *L* instruments. Single-sample individual data MR assumes that for one sample *l*, *Y*_*li*_, *D*_*li*_, *Z*_*li*_ follows a linear structural model common in classical econometrics (Lawlor et al., 2008).

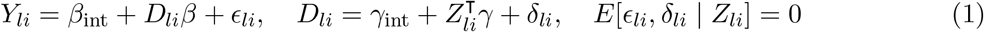

The parameter of interest is *β* and has a causal interpretation under some assumptions (Holland, 1988; Kang et al., 2016; Zhao et al., 2019). Two-sample individual-data MR assumes the same underlying structural model (1) for both samples. But, for sample *l* = 1, the investigator only sees (*Y*_1*i*_, *Z*_1*i*_) and for sample *l* = 2, the investigator only sees (*D*_2*i*_, *Z*_2*i*_) (Pierce and Burgess, 2013; Burgess et al., 2013); this is identical to the setup in Angrist and Krueger (1992). Finally, in two-sample summary-data MR, only summarized statistics of (*Y*_1*i*_, *Z*_1*i*_) and (*D*_2*i*_, *Z*_2*i*_) are available. Specifically, from *n*_1_ samples of (*Y*_1*i*_, *Z*_1*i*_), we obtain (i)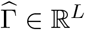 where 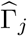 is the estimated association between IV *Z*_1*ij*_ and *Y*_1*i*_ and (ii) 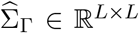, the estimated covariance of 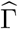. Similarly, from *n*_2_ samples of (*D*_2*i*_, *Z*_2*i*_), we obtain (i) 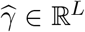 where 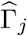 is the estimated association between IV *Z*_2*ij*_ and *D*_2*i*_ and (ii) 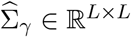, the estimated covariance of 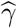. We assume that the summary statistics 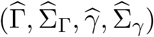 used in the analysis satisfy the following assumptions:

#### Assumption 1.

The IV-exposure effect 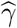 and the IV-outcome effect 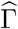 are independent, 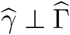.

#### Assumption 2.

The two effect estimates are distributed as 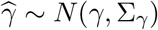 and 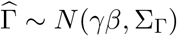

#### Assumption 3.

We have 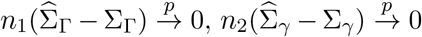 and 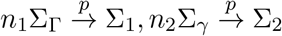 where Σ_1_,Σ_2_ are deterministic positive-definite matrices.

#### Assumption 4.

For some constant *C* ∈ ℝ^*L*^, we have 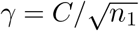

Assumption 1 is typically satisfied in MR by having two GWAS that independently measure SNPs’ associations to the exposure and outcome (Pierce and Burgess, 2013). Assumption 2 is reasonable in publicly-available GWAS where the effect estimates are based on running ordinary least square (OLS) between each instrument and exposure/outcome and since the sample size for each GWAS is on the order of hundreds or thousands, the normality of OLS estimates is plausible. Assumption 3 states that the estimated standard errors converge to their asymptotic variances. Assumption 3 is plausible since the covariance matrices are estimated from OLS residual errors and most of MR assumes that the SNPs are independent of each other. Overall, Assumptions 1–3 are common in two-sample summary-data MR (Zhao et al., 2018). Finally, Assumption 4 is also known as weak-IV asymptotics (Staiger and Stock, 1997) and it provides an asymptotic framework to study the behavior of IV estimators when instruments are weak.

We remark that the literature also assume the instruments are independent to each other, which we do not explicitly impose here. Also, to focus on our contributions to weak IVs in two-sample summary-data MR, we assume that the instruments are valid (e.g. the exclusion restriction holds).

### 2.2 Weak-IV Robust Tests for the Exposure Effect *β*

Consider the null hypothesis *H*_0_ : *β* = *β*_0_ and the alternative *H*_*a*_ : *β* ≠ *β*_0_. We define two statistics *S*(*β*_0_) ∈ ℝ^*L*^ and *R*(*β*_0_) ∈ ℝ^*L*^ from the summary statistics 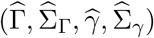.

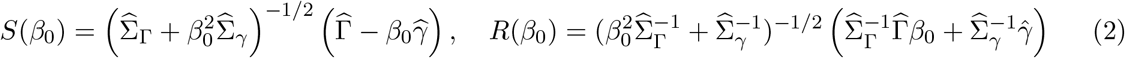

The statistics *S*(*β*_0_) and *R*(*β*_0_) are similar to the independent sufficient statistics of *β* and *π* in the traditional econometric setting (i.e. one-sample individual-data setting) (Moreira, 2003) or in the two-sample individual data setting (Choi et al., 2018). A key difference is that (2) are computed with two-sample summary data. While not exactly sufficient for *β* and *π* in our setting, *S*(*β*_0_) and *R*(*β*_0_) is asymptotically independent as the following Lemma shows.

#### Lemma 1.

If Assumptions 1–4 hold and *n*_1_/*n*_2_ → *c* ∈ (0, ∞), 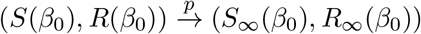
where

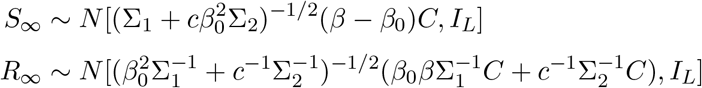

and *S*_∞_ and *R*_∞_ are independent.

The asymptotic independence is crucial as it allows us to follow Moreira (2003) and Andrews et al. (2006) and use *S*(*β*_0_) and *R*(*β*_0_) to construct AR, K, and CLR tests for two-sample summary-data MR.

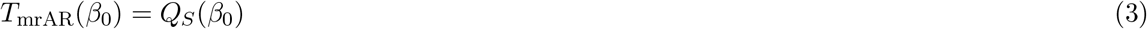

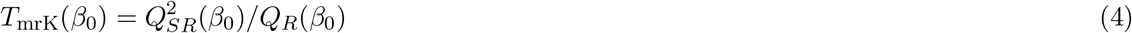

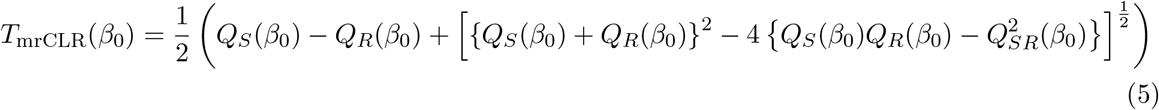

Here, *Q*_*S*_(*β*_0_) = *S*^T^(*β*_0_)*S*(*β*_0_), *Q*_*SR*_ = *S*^T^(*β*_0_)*R*(*β*_0_), and *Q*_*R*_ = *R*^T^(*β*_0_)*R*(*β*_0_). Suppose 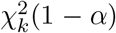 is the 1 − *α* quantile of a Chi-square distribution with *k* degrees of freedom and 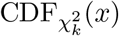 is the cumulative distribution function of a Chi-square distribution with *k* degrees of freedom. The following theorem shows that *T*_mrAR_(*β*_0_), *T*_mrK_(*β*_0_), and *T*_mrCLR_(*β*_0_) have asymptotically pivotal distributions under the null *H*_0_ : *β* = *β*_0_.

#### Theorem 1.

*Suppose Assumptions 1–4 and H*_0_ : *β* = *β*_0_ *hold. For any α* ∈ (0, 1), *as n*_1_, *n*_2_ → ∞, *n*_1_/*n*_2_ → *c* ∈ (0, ∞), *we have*

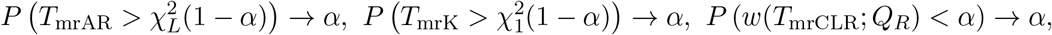

*where*

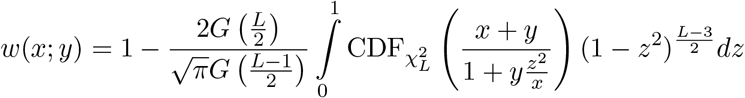

*and G*(·) *is the gamma function.*

Theorem 1 shows that under *H*_0_ : *β* = *β*_0_, the two-sample summary data versions of the AR, K, and CLR tests converge to the classical null distributions for the three tests under the single-sample individual data setting. In particular, like the original CLR test, mrCLR test requires solving the integral *w*(*x*; *y*) to obtain critical values; this integral can be computed by using off-the-shelf numerical integral solvers. We can also use the duality between tests and confidence intervals to derive asymptotically valid 1 − *α* confidence intervals for each test.

In the supplementary materials, we extend these results and construct a point estimator based on minimizing the mrAR test statistic. We show that when the estimated covariance matrices 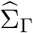 and 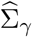 are diagonal, 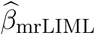 is equivalent to the estimator proposed by Zhao et al. (2018).

## 3 Simulations

We conduct simulation studies to study the performance of our test statistics. The data is generated from the structural model in (1) with *n*_1_ = *n*_2_ = 100,000, on the same order as the sample size in the data analysis section 4. The random error (*δ*_1*i*_, *δ*_2*i*_) is generated from a bivariate standard Normal and the random error *ϵ*_*li*_ is equal to *ϵ*_*li*_ = *ρδ*_*li*_ + (1 − *ρ*^2^)^1/2^*e*_*li*_; the term *e*_*li*_ is from an independent standard Normal and *ρ* = 0.1. We remark that *ρ* signifies the endogeneity between the outcome and the exposure. The *L* = 10 instruments take on values 0, 1, 2, similar to how SNPs are recorded in GWAS, and are generated independently from a Binomial distribution Binom(2, *p*_*j*_), *j* = 1,…, *L* with *p*_*i*_ drawn from a uniform distribution Unif(0.1, 0.9). After generating individual-level data, we compute the summary statistics for sample *l* = 1, 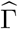 and 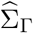, by running an OLS regression between *Y*_1*i*_ and *Z*_1*ij*_ for each instrument *j* and extracting the estimated coefficient and standard error. Similarly, we compute the summary statistics for sample *l* = 2, 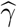 and 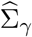 by running an OLS regression between *D*_2*i*_ and *Z*_2*ij*_ for each instrument. The simulation varies the exposure effect *β* and the IV-exposure relationship *γ*. The simulation is repeated 1,000 times.

We examine the power of our proposed tests, *T*_mrAR_, *T*_mrK_, and *T*_mrCLR_ and the power of existing tests in MR, specifically tests based on the IVW estimator, MR-Egger regression, the W.Median estimator, all implemented in the software `Mendelianrandomization` (Yavorska and Burgess, 2017), and MR-RAPs without the robust loss function (Zhao et al., 2018). Figure 1 shows the power curves when the null hypothesis is *H*_0_ : *β* = 0 (left panel) or *H*_0_ : *β* = 1 (right panel); significance level is set at *α* = 0.05. The top panels shows the case when *γ* ranges from {(*r*−0.5)/*n*_1_}^1/2^ to {(*r*+0.5)/*n*_1_}^1/2^ and *r* = 1 the bottom panel shows the case when *r* = 4; the value *r* approximately corresponds to the first-stage F-statistic test typically used to test instrument strength. Under *H*_0_ : *β* = 0, all tests correctly control Type I error under *r* = 1 and *r* = 4. But, our three tests, IVW, and MR-RAPs have power under *r* = 1 and *r* = 4 cases, with *T*_mrCLR_ having the best power among them; this is in agreement with Andrews et al. (2006) who showed that the CLR test in the single-sample individual data setting is nearly optimal. Under *H*_0_ : *β* = 1, none of the pre-existing methods except MR-RAPs have Type I error control when instruments are weak. In contrast, our tests always maintain Type I error control. Also, our tests have power under the alternative, with *T*_mrCLR_ having the best power among them. In fact, in the supplementary materials, we show that tests based on the IVW estimator, weighted median estimator, and MR Egger regression only locally control Type I error at the null *H*_0_ : *β* = 0 when the instruments are weak.

**Figure 1:**
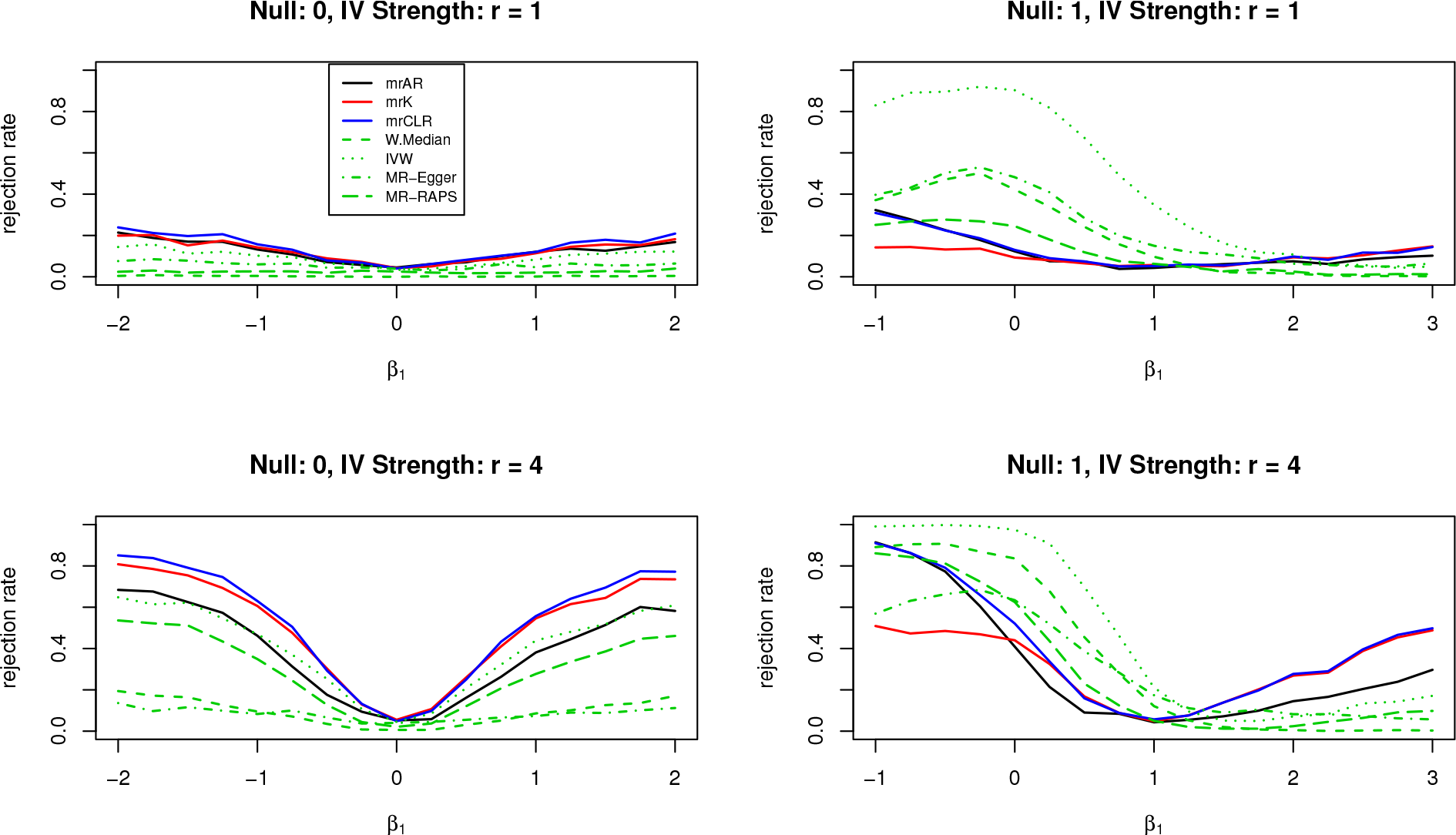
Power curves under different null and IV strength. The left panel is under *H*_0_ : *β*_0_ = 0 and the right panel is under *H*_0_ : *β* = 1. The top panel sets instrument strength to *r* = 1 and the bottom panel sets instrument strength to *r* = 4; *r* approximately corresponds to the first-stage F-statistic test for IV strength.

## 4 Data Analysis

We validate our proposed tests by considering a prototypical MR study on the relationship between BMI and systolic blood pressure where the exposure effect is known to be positive. We use the data set prepared by Zhao et al. (2018) where the authors used three independent GWAS, one from the UK Biobank GWAS (SBP-UKBB) and the other two from GWAS by the Genetic Investigation of ANthropometric Traits (GIANT) consortium (Locke et al., 2015). The BMI-MAL and SBP-UKBB datasets provide summary statistics of the IV-exposure and IV-outcome statistics, respectively. The BMI-FEM dataset is a selection GWAS containing summary statistics of the IV-exposure relationship and is used to pre-screen for strong and uncorrelated IVs.

We conduct two types of analysis with the data. First, we use the data as provided and examine differences between the IVW, weighted median, MR-Egger estimator, MR-RAPS with a robust loss function, and our methods when either *L* = 25 or *L* = 160 instruments are used. The results are in Table 1.

**Table 1:** MR study concerning the effect of body mass index on systolic blood pressure. Parentheses represent 95% confidence intervals.

|  | 25 Instruments | 160 Instruments |
| --- | --- | --- |
| mrK | (0.205, 0.530) | (0.377, 0.771) |
| mrCLR | (0.211, 0.524) | (0.415, 0.731) |
| mrAR | (∅) | (∅) |
| IVW | 0.332 (0.063, 0.600) | 0.317 (0.101, 0.534) |
| W.Median | 0.520 (0.280, 0.759) | 0.522 (0.318, 0.726) |
| MR-Egger | 0.622 (0.101, 1.143) | 0.452 (0.112, 0.792) |
| MR-RAPS | 0.354 (0.097, 0.610) | 0.378 (0.141, 0.615) |

We see that almost all methods provide similar 95% confidence intervals. Even though the weighted median, MR-RAPS, and MR-Egger are robust to invalid instruments, their confidence intervals are similar to confidence intervals from *T*_mrK_ and *T*_mrCLR_, which are not robust to invalid instruments. This suggests either that invalid instruments play a minimal role in this data or, as Small and Rosenbaum (2008) suggests, in the presence of invalid IVs, the first-order bias comes not from invalid IVs, but from weak IVs. This is also based on the observation that mrK produced two split intervals, one in the negative region (−14.375, −10.905) (when *L* = 25) or (−10.376, −6.447) (when *L* = 160) and the other in the positive region. We only report the positive region in Table 1 since we know a priori that the effect is positive; when the exposure effect direction is unknown, we recommend taking the union of the intervals. Surprisingly, *T*_mrAR_ produces an empty interval. This behavior may be an indication that the model is incorrect or that the test lacks power; we plan on investigating this property of *T*_mrAR_ in future work.

Second, inspired by Bound et al. (1995), we “stress-test” the pre-existing methods and replace each of the original IV-exposure effect 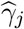 in BMI-MAL and the IV-outcome effect 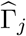 in SBP-UKBB by the values generated below.

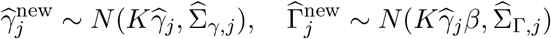

Here, *β* is the true exposure effect and is set to be 0.5 and 1.5. The parameter *K* controls IV strength and ranges from 0 to 1. Under *K* = 1, the new IV-exposure and IV-outcome effects are essentially the original effects, but with a known exposure effect value *β*. But, as *K* decreases to 0, the IV becomes weaker than the original ones. In the extreme case when *K* = 0, there is no way to consistently estimate *β*; the new IV-exposure and IV-outcome effects look statistically indistinguishable if the true exposure effect is *β* = 1 or *β* = 1000.

An ideal confidence interval should be able to (i) simultaneously and automatically detect the lack of identification of the exposure effect *β* by producing an infinite confidence interval when *K* = 0 and a bounded confidence intervals as *K* moves away from zero and (ii) for all values of *K*, provide 95% coverage. As Figure 2 shows, when we run the existing MR methods, none of them achieve these two goals. At *K* = 0, they always produce bounded confidence intervals, even though there is no way to identify the exposure effect from data. Specifically, when *K* = 0, *β* = 0.5, the confidence interval given by Weighted median, IVW, MR-Egger and MR-RAPS are (−0.624, 0.321), (−0.411, 0.265), (−0.527, 0.624), and (−0.460, 0.020), respectively. They are bounded and only the confidence interval generated from MR-Egger covers the true effect *β* = 0.5. But *T*_mrAR_, *T*_mrK_, *T*_mrCLR_ produce unbounded confidence intervals. We observe a similar phenomena when *K* = 0 and *β* = 1.5: the confidence intervals generated from Weighted median, IVW, MR-Egger and MR-RAPS are (−0.385, 0.547), (−0.312, 0.330), (−0.826, 0.317), and (−0.575, 0.942), respectively. All of them are bounded and fail to cover the true causal effect, but our tests produce unbounded confidence intervals. Also, when *K* is near zero so that the IV-exposure is sufficiently weak, all methods except MR-RAPS fail to achieve 95% coverage. In contrast, our tests always satisfy the two criterions (i) and (ii). They automatically produce infinite confidence intervals when *K* = 0 to alert the researcher about lack of identification and produce bounded intervals as *K* moves away from zero. They also always maintain 95% coverage for any value of *K*. In short, our proposed tests adapt to the data and always produce honest intervals regardless of the underlying instrument strength.

**Figure 2:**
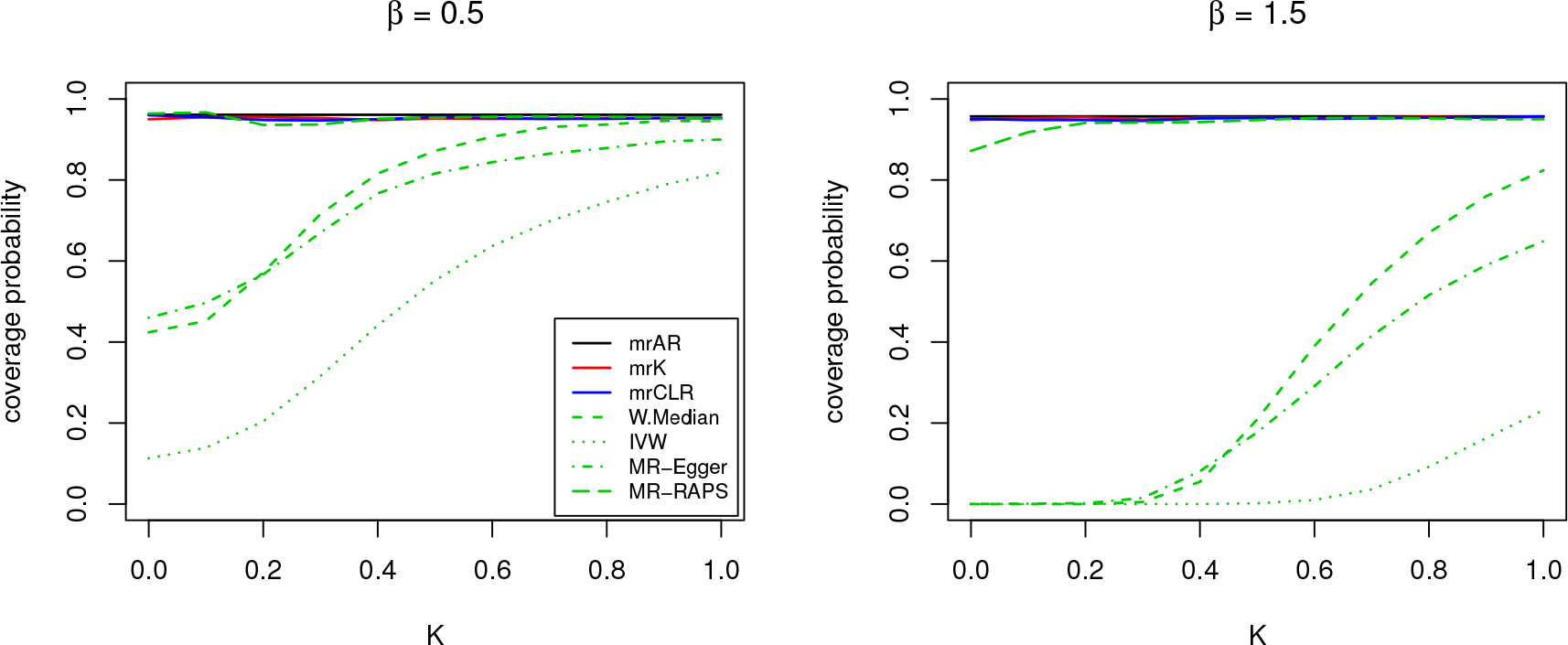
Coverage probability under different IV strength. The left panel sets the true causal effect *β* to be 0.5 and the right panel sets *β* to be 1.5.

## 5 Conclusion

In this paper, we propose weak-IV robust test statistics for two-sample summary data in MR. We extend the existing AR, Kleibergen, and CLR tests in econometrics and show that it has the same Type I error control under weak instrument asymptotics. The numerical results show that the proposed tests, especially the mrCLR test, have better size control and power compared to current methods when the instruments are weak. Additionally, when we stress-test different methods, our methods, especially the CLR test, adapts to the underlying instrument strength and provides valid 95% coverage. In practice, we recommend MR investigators use the mrCLR test to test exposure effects as it provides valid confidence intervals regardless of IV strength. The code to implement our tests is in the supplementary materials

