## supplementary material for "Weak-Instrument Robust Tests in Two-Sample Summary-Data Mendelian Randomization"

### Supplementary Materials for "Weak-Instrument Robust Tests in Two-Sample Summary-Data Mendelian Randomization"

**Abstract**

In this supplementary material for the paper *Weak-Instrument Robust Tests in Two-Sample Summary-Data in Mendelian Randomization*, we propose a set of robust point estimators for the exposure effect and provide additional results for simulation in Section 4 of the main manuscript. We also prove our results in the main paper.

#### 1 Robust Point Estimators for the Exposure Effect $\beta$

In this section, we extend the LIML estimator (Anderson et al., 1949) and the recent unbiased estimator under known signs (Andrews and Armstrong, 2017) to the two-sample summary data setting. We make the same assumptions on the summary statistics as in Section 2 of the main manuscript

**Assumption 1.** The IV-exposure effect $\widehat{\gamma}$ and the IV-outcome effect $\widehat{\Gamma}$ are independent, $\widehat{\gamma} \perp \widehat{\Gamma}$.

**Assumption 2.** The two effect estimates follow $\widehat{\gamma} \sim N(\gamma, \Sigma_\gamma), \widehat{\Gamma} \sim N(\gamma\beta, \Sigma_\Gamma)$

**Assumption 3.** We have $n_1(\widehat{\Sigma}_\Gamma - \Sigma_\Gamma) \xrightarrow{p} 0$, $n_2(\widehat{\Sigma}_\gamma - \Sigma_\gamma) \xrightarrow{p} 0$ and $n_1\Sigma_\Gamma \xrightarrow{p} \Sigma_1, n_2\Sigma_\gamma \xrightarrow{p} \Sigma_2$, where $\Sigma_1, \Sigma_2$ are deterministic positive-definite matrices.

**Assumption 4.** For some constant $C \in \mathbb{R}^L$, we have $\gamma = C/\sqrt{n_1}$

In the single-sample individual-data setting, the limited information maximum likelihood(LIML) estimator is a likelihood-based estimator of $\beta$ by Anderson et al. (1949) that exhibits robustness under many weak instruments (Stock and Yogo, 2005; Chao and Swanson, 2005) where in addition to Assumption 4, the number of instruments is allowed to grow at a certain rate. Additionally, in the one-sample individual-data setting, the LIML estimator is equivalent to the minimum of the Anderson-Rubin test (Stock et al., 2002).

Following the construction of the original LIML, we also propose a point estimator for $\beta$, which we denote it as $\widehat{\beta}_{\mathrm{mrLIML}}$ as the solution to the minimum of the two-sample summary data AR test $T_{\mathrm{mrAR}}(\beta_0)$, i.e.

$$\widehat{\beta}_{\mathrm{mrLIML}} := \mathrm{argmin}_{\beta} T_{\mathrm{mrAR}}(\beta_0) \tag{1}$$

When the estimated covariance matrices $\widehat{\Sigma}_{\Gamma}$ and $\widehat{\Sigma}_{\gamma}$ of the effect estimates are diagonal, $\widehat{\beta}_{\mathrm{mrLIML}}$ is equivalent to the estimator proposed by Zhao et al. (2018). But, our formulation can handle non-diagonal covariance matrices, i.e. when either the instruments or the marginal effects are correlated. To evaluate $\widehat{\beta}_{\mathrm{mrLIML}}$, we can (i) do a simple grid search since $\beta$ is a scalar parameter or (ii) gradient descent where the local minimum can be found by iteratively taking steps proportional to the negative of first-order derivative of $T_{\mathrm{mrAR}}(\beta_0)$.

The next estimator is based on an assumption that the sign of the effect of the instrument on the exposure is known a prioiri. For example, in an MR study by Kang et al. (2016) concerning the study of the effect of malaria (i.e. exposure) on stunted growth (i.e. outcome) in children in sub-Saharan Africa, the authors used a gene, known as the sickle cell trait, as an instrument. It was well-known biologically that carrying a copy of this trait reduced malarial infections, or more precisely, the sign of the IV-exposure $\gamma$ was known a priori. A recent work by Andrews and Armstrong (2017) proposed an unbiased estimator of the exposure effect $\beta$ provided that the sign of the IV-exposure is known and the standard errors of the estimates are known. Unfortunately, their work was limited to the one-sample individual-data setting.

We build on this recent work and the work by Voinov and Nikulin (2012), where the latter proposed an unbiased estimator for the inverse of the mean parameter in Normally distributed data, and propose a two-sample summary data variant of the unbiased estimator. Let $\mathrm{CDF}_{\mathrm{N}}(x)$ and $\mathrm{PDF}_{\mathrm{N}}(x)$ denote the cumulative distribution function and the probability density function, respectively, of the standard Normal distribution. For two-sample summary data setting and each IV $j$, consider the following estimator $\beta$

$$\widehat{\beta}_{j,\mathrm{U}} = \frac{\widehat{\Gamma}_j}{\sqrt{\Sigma_{jj,\gamma}}} \cdot \frac{1 - \mathrm{CDF}_{\mathrm{N}}\left(\frac{\widehat{\gamma}_j}{\sqrt{\Sigma_{jj,\gamma}}}\right)}{\mathrm{PDF}_{\mathrm{N}}\left(\frac{\widehat{\gamma}_j}{\sqrt{\Sigma_{jj,\gamma}}}\right)}, \tag{2}$$

where $\Sigma_{jj,\gamma}$ is the $j$ diagonal element of $\Sigma_{\gamma}$. Theorem 1 shows that the estimator is unbiased when the sign of $\gamma_j$ and the standard error of $\widehat{\gamma}_j$ are known.

**Theorem 1.** *Suppose assumptions 1 and 2 hold where, without loss of generality, $\gamma_j > 0$. Then if $\Sigma_{jj,\gamma}$ is known, $\widehat{\beta}_{j,\mathrm{U}}$ in equation (2) is an unbiased estimator of $\beta$.*

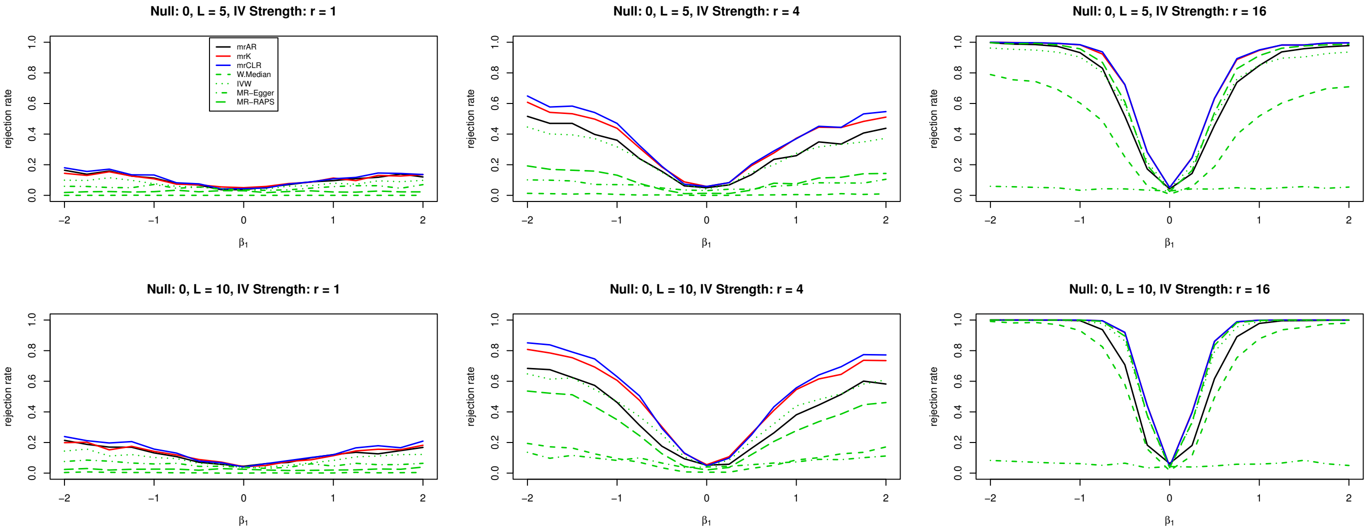


Figure 1: Power Curves under Different number of IVs and IV Strength. Null: 0. The top panel sets the number of instrument to $L = 5$ and the bottom panel sets the number of instruments to $L = 10$; The left panel sets instrument strength to $r = 1$, the middle panel sets instrument strength to $r = 4$ and the right panel sets instrument strength to $r = 16$; $r$ approximately corresponds to the first-stage F-statistic test for IV strength.

Applying Theorem 1 in practice requires plugging in an estimator of the standard error of $\Sigma_{jj,\gamma}$ and with modern MR involving tens of thousands samples, $\Sigma_{jj,\gamma}$ can be accurately estimated. Also, because Theorem 1 holds for each instrument $j$, when multiple instruments are available, we can aggregate $\widehat{\beta}_{j,\mathrm{U}}$s to obtain a single unbiased estimator. For example, any linear combination of $\widehat{\beta}_{j,\mathrm{U}}$ is still unbiased for $\beta$. In our numerical works, we opt to use the simple average to aggregate our $L$ unbiased estimates, $\widehat{\beta}_{\mathrm{U}} = \frac{1}{L}\sum_{j=1}^{L}\widehat{\beta}_{j,\mathrm{U}}$

#### 2 Extended Simulation Results

In this section, we provide extended simulation results to Section 3 of the main manuscript. The data generating process is the same as the main manuscript.

We first examine the performance of our proposed tests $T_{\mathrm{mrAR}}, T_{\mathrm{mrK}}$, and $T_{\mathrm{mrCLR}}$, as well as existing tests in MR, specifically tests based on the IVW estimator, MR-Egger regression, the W.Median estimator, all implemented in the software `Mendelianrandomization` (Yavorska and Burgess, 2017), and MR-RAPs without the robust loss function (Zhao et al., 2018). Figure 1 shows the power curves under the null hypothesis $H_0 : \beta = 0$. Each graph uses a different pair of $(L, r)$. The top panel shows the case when $L = 5$ and the bottom panel $L = 10$. The IV strength is set as $r = 1, 4, 16$ respectively in the left, middle and right panel. All the methods correctly control Type I error. But, our three tests, IVW, and MR-RAPS have power under $r = 1$ and $r = 4$ (when the

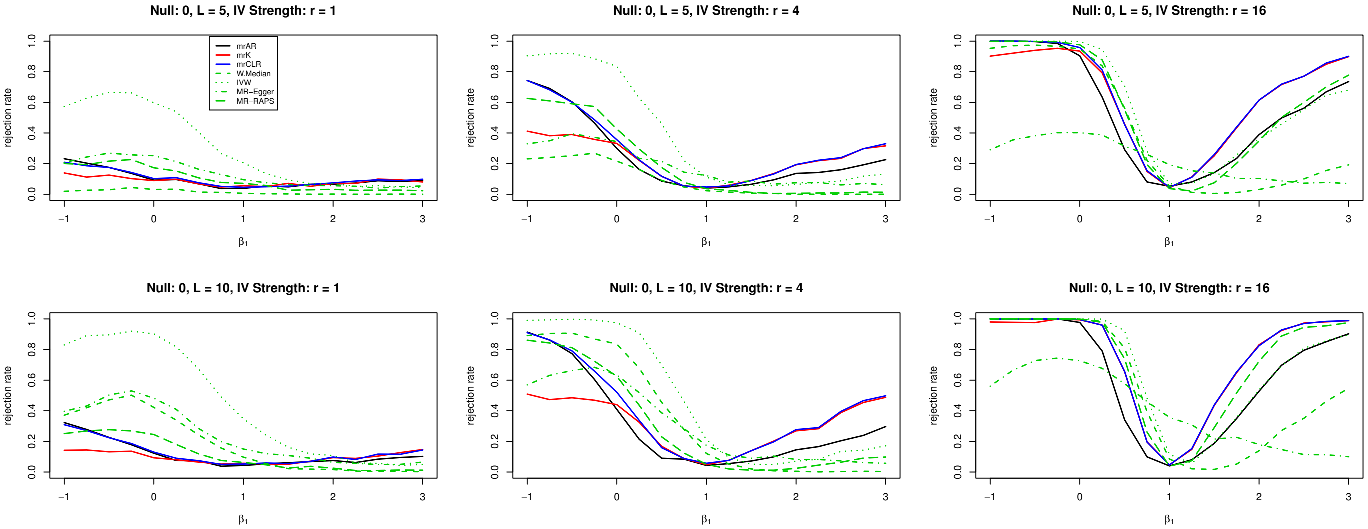


Figure 2: Power Curves under Different number of IVs and IV Strength. Null: 1. The top panel sets the number of instrument to $L = 5$ and the bottom panel sets the number of instruments to $L = 10$; The left panel sets instrument strength to $r = 1$, the middle panel sets instrument strength to $r = 4$ and the right panel sets instrument strength to $r = 16$; $r$ approximately corresponds to the first-stage F-statistic test for IV strength.

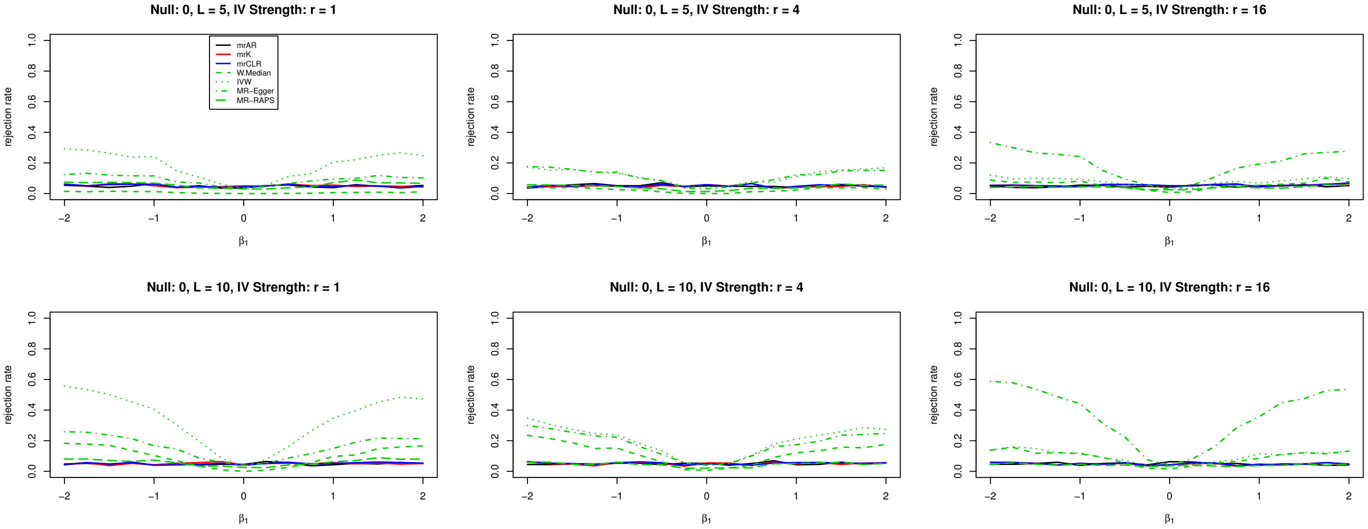


Figure 3: Size Curves under Different number of IVs and IV Strength. The top panel sets the number of instrument to $L = 5$ and the bottom panel sets the number of instruments to $L = 10$; The left panel sets instrument strength to $r = 1$, the middle panel sets instrument strength to $r = 4$ and the right panel sets instrument strength to $r = 16$; $r$ approximately corresponds to the first-stage F-statistic test for IV strength.

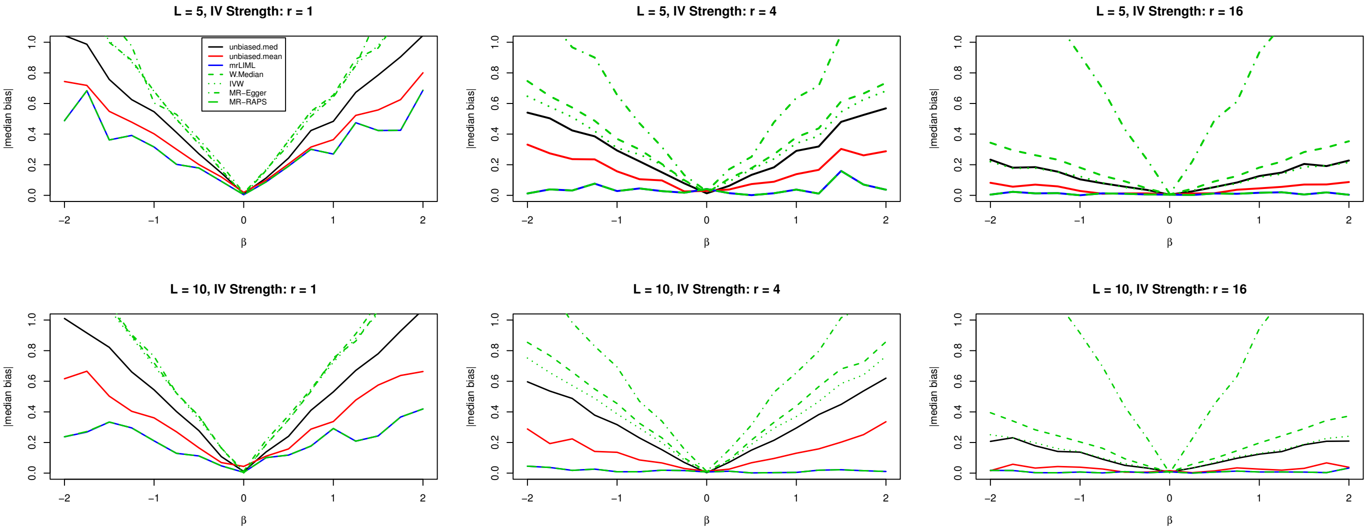


Figure 4: Absolute Median Bias of Point Estimators. The top panel sets the number of instrument to $L = 5$ and the bottom panel sets the number of instruments to $L = 10$; The left panel sets instrument strength to $r = 1$, the middle panel sets instrument strength to $r = 4$ and the right panel sets instrument strength to $r = 16$; $r$ approximately corresponds to the first-stage F-statistic test for IV strength.

instruments are weak), with $T_{\mathrm{mrCLR}}$ having the best power among them; this is in agreement with Andrews et al. (2006) who showed that the CLR test in the single-sample individual data setting is nearly optimal. When the instruments are relatively stronger ($r = 16$), all methods except MR-Egger have power and $T_{\mathrm{mrCLR}}$ has the best power among them. Figure 2 shows the power curves under $H_0 : \beta = 1$. In this scenario, none of the pre-existing methods except MR-RAPs have Type I error control when instruments are weak while our tests always maintain Type I error control. Also, our tests have power under the alternative, with $T_{\mathrm{mrCLR}}$ having the best power among them. Figure 3 shows the Type I error under $H_0 : \beta = \beta_0$ with $\beta_0$ ranging from $-2$ to 2. All the tests are carried out under the null. The size distortion of all the pre-existing methods increases as the true exposure effect $\beta$ moves away from 0, but our tests always have size control. Overall, we find that our proposed tests are superior to pre-existing MR methods, especially under weak instruments, with respect to Type I error control and power and $T_{\mathrm{mrCLR}}$ has the best power in all simulated scenarios.

Second, we examine the performance of our point estimators, $\widehat{\beta}_{\mathrm{mrLIML}}$ and the $\widehat{\beta}_U$ and compare it against the same set of pre-existing MR methods; for the unbiased estimator, we assume that the sign of each IV-exposure effect is positive. The simulation setting is the same as before, except we vary the true exposure effect $\beta$ from $-2$ to 2. We measure the absolute median bias and the median absolute deviation (MAD) of all the estimators and the results are shown in Figure 4 and Figure 5. In terms of absolute median bias, our LIML estimator and unbiased estimator are less biased than pre-existing methods, with the notable exception to MR-RAPS; we also find that LIML is the

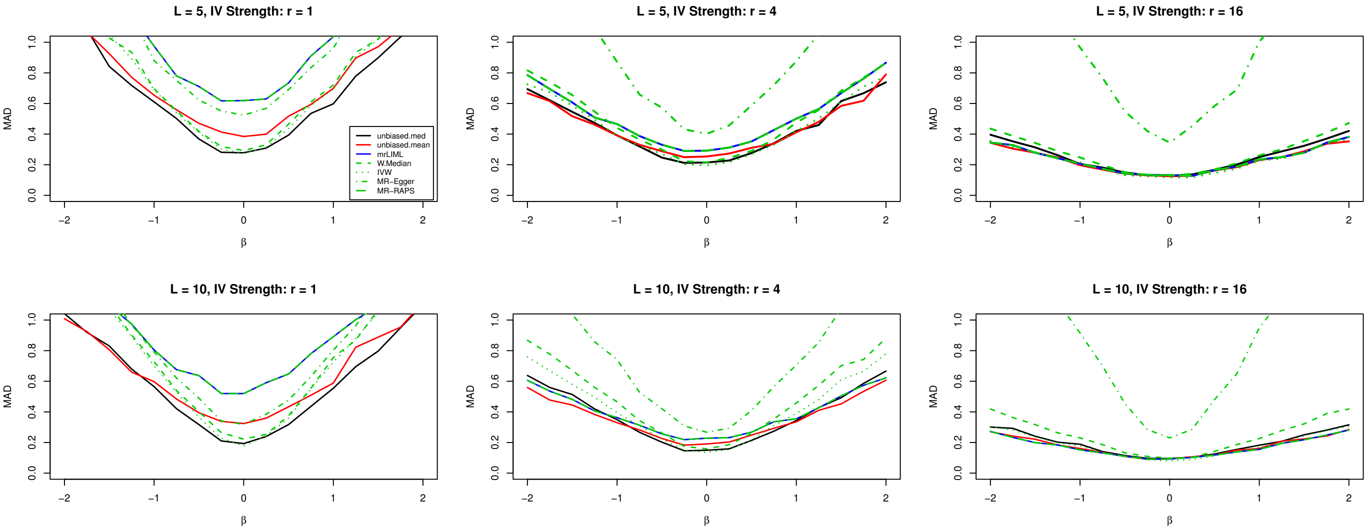


Figure 5: MAD of Point Estimators. The top panel sets the number of instrument to $L = 5$ and the bottom panel sets the number of instruments to $L = 10$; The left panel sets instrument strength to $r = 1$, the middle panel sets instrument strength to $r = 4$ and the right panel sets instrument strength to $r = 16$; $r$ approximately corresponds to the first-stage F-statistic test for IV strength.

least biased overall. Also, similar to the testing case, as the true exposure effect moves away from zero, the IVW estimator, the weighted median estimator , MR-Egger estimator, and the unbiased estimator are biased; in contrast, LIML is unaffected by the value of $\beta$. With respect to median absolute deviation, all methods are similar around 0. But with larger $|\beta|$, the MAD of the LIML, MR-RAPs, and the unbiased estimator are smaller than the IVW, weighted median, and MR-Egger estimators.

### 3 Proof of Lemmas and Theorems

*Proof of Lemma 1.*

Define

$$\tilde{S}(\beta_0) = \left(\Sigma_\Gamma + \beta_0^2 \Sigma_\gamma\right)^{-1/2} \left(\widehat{\Gamma} - \beta_0 \widehat{\gamma}\right),$$

$$\tilde{R}(\beta_0) = (\beta_0^2 \Sigma_\Gamma^{-1} + \Sigma_\gamma^{-1})^{-1/2} \left(\Sigma_\Gamma^{-1} \widehat{\Gamma} \beta_0 + \Sigma_\gamma^{-1} \hat{\gamma}\right)$$

Let $d_1 = \hat{\Gamma} - \Gamma, d_2 = \hat{\gamma} - \gamma$, then we have

$$\begin{aligned}
\tilde{S}(\beta_0) &= (\Sigma_\Gamma + \beta_0^2 \Sigma_\gamma)^{-1/2} [\hat{\Gamma}, \hat{\gamma}] b_0 \\
&= (n_1 \Sigma_\Gamma + n_1 \beta_0^2 \Sigma_\gamma)^{-1/2} n_1^{1/2} (I_L, -\beta_0 I_L)[vec(\Gamma, \gamma) + vec(d_1, d_2)] \\
&\xrightarrow{p} (\Sigma_1 + c\beta_0^2 \Sigma_2)^{-1/2} (\beta - \beta_0) C + (\Sigma_\Gamma + \beta_0^2 \Sigma_\gamma)^{-1/2} (I_L, -\beta_0 I_L) vec(d_1, d_2) \equiv S_\infty \\
&\sim N((\Sigma_1 + c\beta_0^2 \Sigma_2)^{-1/2} (\beta - \beta_0) C, I_L)
\end{aligned}$$

$$\begin{aligned}
\tilde{R}(\beta_0) &= (\beta_0^2 \Sigma_\Gamma^{-1} + \Sigma_\gamma^{-1})^{-1/2} (\Sigma_\Gamma^{-1} \beta_0, \Sigma_\gamma^{-1}) vec(\hat{\Gamma}, \hat{\gamma}) \\
&= (\beta_0^2 n_1^{-1} \Sigma_\Gamma^{-1} + n_1^{-1} \Sigma_\gamma^{-1})^{-1/2} [n_1^{-1} \Sigma_\Gamma^{-1} \beta_0, n_1^{-1} \Sigma_\gamma^{-1}] n_1^{1/2} vec(\Gamma, \gamma) + (\beta_0^2 \Sigma_\Gamma^{-1} + \Sigma_\gamma^{-1})^{-1/2} [\Sigma_\Gamma^{-1} \beta_0, \Sigma_\gamma^{-1}] \\
&\quad vec(d_1, d_2) \\
&\xrightarrow{p} (\beta_0^2 \Sigma_1^{-1} + c^{-1} \Sigma_2^{-1})^{-1/2} [\beta_0 \beta \Sigma_1^{-1} C + c^{-1} \Sigma_2^{-1} C] + (\beta_0^2 \Sigma_{n,\Gamma}^{-1} + \Sigma_{n,\gamma}^{-1})^{-1/2} (\Sigma_{n,\Gamma}^{-1} \beta_0, \Sigma_{n,\gamma}^{-1}) vec(d_1, d_2) \equiv R_\infty \\
&\sim N((\beta_0^2 \Sigma_1^{-1} + c^{-1} \Sigma_2^{-1})^{-1/2} (\beta_0 \beta \Sigma_1^{-1} C + c^{-1} \Sigma_2^{-1} C), I_L)
\end{aligned}$$

Since $n_1(\hat{\Sigma}_\Gamma - \Sigma_\Gamma) \xrightarrow{p} 0, n_2(\hat{\Sigma}_\gamma - \Sigma_\gamma) \xrightarrow{p} 0$, we have that $((S(\beta_0), R(\beta_0)) - (S_\infty, R_\infty)) \xrightarrow{p} 0$. The asymptotic normal distributions of $S_n$ and $R_n$ are independent because they are non-stochastic functions of $(\Sigma_\Gamma + \beta_0^2 \Sigma_\gamma)^{-1/2} (I_L, -\beta_0 I_L) vec(d_1, d_2)$ and $(\beta_0^2 \Sigma_\Gamma^{-1} + \Sigma_\gamma^{-1})^{-1/2} (\Sigma_\Gamma^{-1} \beta_0, \Sigma_\gamma^{-1}) vec(d_1, d_2)$, and

$$\begin{aligned}
&Cov((\Sigma_\Gamma + \beta_0^2 \Sigma_\gamma)^{-1/2} (I_L', -\beta_0 I_L') vec(d_1, d_2), (\beta_0^2 \Sigma_\Gamma^{-1} + \Sigma_\gamma^{-1})^{-1/2} (\Sigma_\Gamma^{-1} \beta_0, \Sigma_\gamma^{-1}) vec(d_1, d_2)) \\
&= (\Sigma_\Gamma + \beta_0^2 \Sigma_\gamma)^{-1/2} (I_L, -\beta_0 I_L) \begin{pmatrix} \Sigma_\Gamma & 0 \\ 0 & \Sigma_\gamma \end{pmatrix} (\Sigma_\Gamma^{-1} \beta_0, \Sigma_\gamma^{-1})' (\beta_0^2 \Sigma_\Gamma^{-1} + \Sigma_\gamma^{-1})^{-1/2} \\
&= 0.
\end{aligned}$$

*Proof of Theorem 1.*

Let $Q_\infty = \begin{pmatrix} Q_{\infty,S} & Q_{\infty,RS} \\ Q_{\infty,SR} & Q_{\infty,R} \end{pmatrix} = \begin{pmatrix} S_\infty' S_\infty & R_\infty' S_\infty \\ S_\infty' R_\infty & R_\infty' R_\infty \end{pmatrix}, S_2 = Q_{\infty,SR}/(Q_{\infty,S} Q_{\infty,R})^{1/2}$.

By lemma 1, $Q_{\infty,S}$ follows $\chi_L^2$ under $H_0$. Since $T_{\text{mrAR}} \xrightarrow{p} Q_{\infty,S}$, we have that $P(T_{\text{mrAR}} > \chi_L^2(1-\alpha)) \to \alpha$. Similarly, since $Q_{\infty,SR}^2/Q_{\infty,R}$ follows $\chi_1^2$ under $H_0$, and $T_{\text{mrK}} \xrightarrow{p} Q_{\infty,SR}^2/Q_{\infty,R}$, we have that $P(T_{\text{mrK}} > \chi_1^2(1-\alpha)) \to \alpha$.

For $T_{\text{mrCLR}} = \frac{1}{2}(Q_S - Q_R + ((Q_S + Q_R)^2 - 4(Q_S Q_R - Q_{SR}^2))^{1/2})$, let $LR_\infty = \frac{1}{2}(Q_{\infty,S} - Q_{\infty,R} + ((Q_{\infty,S} + Q_{\infty,R})^2 - 4(Q_{\infty,S} Q_{\infty,R} - Q_{\infty,SR}^2))^{1/2})$. By Lemma 1, $T_{\text{mrCLR}} \xrightarrow{p} LR_\infty$.

Let $w(x; q_r) = P_0(LR_\infty > x | Q_{\infty,R} = q_r) = 1 - \frac{2G(\frac{L}{2})}{\sqrt{\pi} G(\frac{L-1}{2})} \int_0^1 \text{CDF}_{\chi_L^2}\left(\frac{x + q_r}{1 + q_r \frac{z^2}{x}}\right)(1 - z^2)^{\frac{L-3}{2}} dz$, where $P_0$ denotes the probability evaluated under $H_0$. We reject $H_0$ if $w(T_{\text{mrCLR}}; Q_R)$ is smaller than $\alpha$. Our goal is to show that $\lim_{n_1,n_2\to\infty} P(w(T_{\text{mrCLR}};Q_R)<\alpha)=\alpha$ when $H_0$ is true.

Denote $F_{0,q_r}(LR_\infty)=w(LR_\infty,q_r)$ as the conditional CDF of $LR_\infty$ given $Q_{\infty,R}=q_r$. Under $H_0$, we have

$$\begin{aligned}
\lim_{n_1,n_2\to\infty} P(w(T_{\text{mrCLR}};Q_R)<\alpha|Q_{\infty,R}=q_r) &= \lim_{n_1,n_2\to\infty} P_0(w(T_{\text{mrCLR}};q_r)<\alpha|Q_{\infty,R}=q_r)\\
&= \lim_{n_1,n_2\to\infty} P_0(F_{0,q_r}(T_{\text{mrCLR}})<\alpha|Q_{\infty,R}=q_r)\\
&= P_0(F_{0,q_r}(LR_\infty)<\alpha|Q_{\infty,R}=q_r)=\alpha\\
\lim_{n_1,n_2\to\infty} P_0(w(T_{\text{mrCLR}};Q_R)>\alpha) &= \lim_{n_1,n_2\to\infty}\int P_0(w(T_{\text{mrCLR}};Q_R)>\alpha|Q_{\infty,R}=q_r)f_{Q_{\infty,R}}(q_r)dq_r\\
&= \int \lim_{n_1,n_2\to\infty} P_0(p-value>\alpha|Q_{\infty,R}=q_r)f_{Q_{\infty,R}}(q_r)dq_r\\
&= \int (1-\alpha)f_{Q_{\infty,R}}(q_r))dq_r = 1-\alpha
\end{aligned}$$

where the exchange of the limit and integral in the second last equality is guaranteed by Dominated Convergence Theorem.

*Proof of Theorem S.1.*

Let $\hat\tau(\widehat\gamma_j,\Sigma_{jj,\gamma})=\frac{1}{\sqrt{\Sigma_{jj,\gamma}}}\cdot\frac{1-\text{CDF}_{\text{N}}\left(\frac{\widehat\gamma_j}{\sqrt{\Sigma_{jj,\gamma}}}\right)}{\text{PDF}_{\text{N}}\left(\frac{\widehat\gamma_j}{\sqrt{\Sigma_{jj,\gamma}}}\right)}$. Since $\frac{\widehat\gamma_j}{\sqrt{\Sigma_{jj,\gamma}}}\sim N(\frac{\gamma}{\sqrt{\Sigma_{jj,\gamma}}},1)$, we have

$$\begin{aligned}
E(\hat\tau(\widehat\gamma_j,\Sigma_{jj,\gamma})) &= \frac{1}{\sqrt{\Sigma_{jj,\gamma}}}E\frac{1-\text{CDF}_{\text{N}}(\widehat\gamma_j/\sqrt{\Sigma_{jj,\gamma}})}{\text{PDF}_{\text{N}}(\widehat\gamma_j/\sqrt{\Sigma_{jj,\gamma}})}\\
&= \frac{1}{\sqrt{\Sigma_{jj,\gamma}}}\int\frac{1-\text{CDF}_{\text{N}}(x)}{\text{PDF}_{\text{N}}(x)}\text{PDF}_{\text{N}}(x-\frac{\gamma}{\sqrt{\Sigma_{jj,\gamma}}})dx\\
&= \frac{1}{\sqrt{\Sigma_{jj,\gamma}}}\int(1-\text{CDF}_{\text{N}}(x))\exp((\gamma_1/\sqrt{\Sigma_{jj,\gamma}})x-(\gamma_1/\sqrt{\Sigma_{jj,\gamma}})^2/2)dx\\
&= \frac{1}{\gamma_1}\exp(-(\gamma_1/\sqrt{\Sigma_{jj,\gamma}})^2/2)\int(1-\text{CDF}_{\text{N}}(x))d(\exp((\gamma_1/\sqrt{\Sigma_{jj,\gamma}})x))\\
&= \frac{1}{\gamma_1}\exp(-(\gamma_1/\sqrt{\Sigma_{jj,\gamma}})^2/2)\{[(1-\text{CDF}_{\text{N}}(x))\exp(\gamma_1/\sqrt{\Sigma_{jj,\gamma}})x]_{x=-\infty}^{\infty}\\
&\quad+\exp((\gamma_1/\sqrt{\Sigma_{jj,\gamma}})x)\text{PDF}_{\text{N}}(x)dx\}\\
&= \frac{1}{\gamma_1}\exp(-(\gamma_1/\sqrt{\Sigma_{jj,\gamma}})^2/2)\int\exp((\gamma_1/\sqrt{\Sigma_{jj,\gamma}})x)\frac{1}{\sqrt{2\pi}}\exp(-\frac{x^2}{2})dx\\
&= \frac{1}{\gamma_1}\int\frac{1}{\sqrt{2\pi}}\exp(-\frac{(x-\gamma_1/\sqrt{\Sigma_{jj,\gamma}})^2}{2})dx\\
&= \frac{1}{\gamma_1}
\end{aligned}$$

Thus $E(\widehat\beta_{j,\text{U}})=E(\widehat\Gamma_j\hat\tau(\widehat\gamma_j,\Sigma_{jj,\gamma}))=\frac{\Gamma_j}{\gamma_j}$ by independence between $\widehat\Gamma_j$ and $\widehat\gamma_j$.
